# Single molecule microscopy to profile the effect of zinc status of transcription factor dynamics

**DOI:** 10.1101/2022.05.10.491421

**Authors:** Leah J. Damon, Jesse Aaron, Amy E. Palmer

## Abstract

Transcription factors (TFs) are DNA binding proteins that control the expression of genes. The regulation of transcription is a complex process that involves binding of TFs to specific sequences, recruitment of cofactors and chromatin remodelers, assembly of the pre-initiation complex and ultimately the recruitment of RNA polymerase II. Increasing evidence suggests that TFs are highly dynamic and interact only transiently with DNA. Single molecule microscopy techniques are powerful approaches for visualizing and tracking individual TF molecules as they diffuse in the nucleus and interact with DNA. In this work, we employ multifocus microscopy and highly inclined and laminated optical sheet microscopy to track TF dynamics in response to perturbations in labile zinc inside cells. We sought to define whether zinc-dependent TFs sense changes in the labile zinc pool by determining whether their dynamics and DNA binding can be modulated by zinc. While it is widely appreciated that TFs need zinc to bind DNA, whether zinc occupancy and hence TF function are sensitive to *changes* in cellular zinc remain open questions. We utilized fluorescently tagged versions of the glucocorticoid receptor (GR), with two C4 zinc finger domains, and CCCTC-binding factor (CTCF), with eleven C2H2 zinc finger domains. We found that the biophysical dynamics of both TFs are susceptible to changes in zinc, but in subtly different ways. These results indicate that at least some transcription factors are sensitive to zinc dynamics, revealing a potential new layer of transcriptional regulation.

## Introduction

Transcription factors (TFs) regulate gene expression through a tightly organized series of events, ultimately leading to the production of RNA by RNA Polymerase (RNAP). While all transcription requires a core group of general TFs to initiate transcription and guide RNAP to its target, many genes require additional cell-type specific TFs that bind to regions of DNA known as enhancer regions around the target gene to further stimulate or even repress gene expression^1,2^. The activities of these TFs are tightly regulated by the cell. TFs tend to remain inactive until the appropriate signal is transduced to them, leading to activation by means such as post-translational modification (e.g., phosphorylation) or translocation from one part of the cell, such as the cytosol, to the nucleus^3,4^. While much of the early work probing TF DNA binding properties was done *in vitro*, many have sought to move towards more *in situ* methods. This has become more feasible with the advent of next-generation sequencing techniques. Chromatin immunoprecipitation with sequencing (ChIP-seq) reveals with relatively good accuracy the sequences at which a given TF is bound with relatively good accuracy. Additionally, computational techniques have been developed that use chromatin accessibility data from ATAC-seq to infer TF binding within regions of open chromatin^5,6^. However, these techniques have their drawbacks. Beyond the requirement of knowing how to prepare high-quality sequencing libraries and how to interrogate the sequencing data, these techniques only provide snapshots of what a particular TF is doing at a given treatment time. Additionally, most genomics assays are bulk assays that report averages from thousands of cells, and it has been widely reported that cell-to-cell heterogeneity can be critical to understanding diseases such as cancer.^7,8^

Microscopy, namely single molecule (SM) microscopy, fills the single cell and single TF niche where genomics assays lag. While a barrier to studying TF activity previously was access to and expression of a fluorescent variant of the TF of interest, this has become much easier of late with the advent of overexpression systems such as the PiggyBac transposon system^9^ and the genome editing technique CRISPR^10,11^. Additionally, the development of robust protein labeling systems, such as HaloTag and its corresponding ligands^12,13^, has allowed researchers to expand into the SM field to monitor TFs in live cells. SM microscopy allows for evaluation of diffusion coefficients, search mechanisms, and dwell (binding) times for individual TFs, revealing intracellular and even intercellular differences amongst populations of TFs. This can also be paired with other imaging tools, such as those that study RNA production^14,15^, to further interrogate the functional consequences of TF activities.

Zinc (Zn^2+^) finger TFs are among the most ubiquitous TF families in human cells, with nearly half (868/1792) of all predicted TFs utilizing Zn^2+^ fingers to interact with genomic targets^16^. Although the precise coordination of the Zn^2+^ ion can vary, all of these share the common attribute that the binding of a Zn^2+^ ion to the protein domain allows for formation of the proper fold that then interacts with DNA^17^. It is well established that *in vitro* Zn^2+^ finger TFs require their Zn^2+^ cofactor to bind DNA, but whether these TFs are sensitive to changes in the Zn^2+^ pool in cells has not been examined, with one exception: the metal-responsive transcription factor (MTF1). MTF1 senses high Zn^2+^ through its array of six Zn^2+^ fingers and in the presence of high Zn^2+^, translocates to the nucleus to regulate metal buffering proteins (metallothionines) Zn^2+^ and metal transporters. While the apparent dissociation constant (K_D_) for Zn^2+^ in the full length MTF1 protein was found to be 31 pM^18^, biochemical studies^19,20^ have shown that Zn^2+^ fingers five and six are the most reactive and have the weakest affinity for cobalt (which is often used to measure affinities of Zn^2+^ binding proteins^17^), suggesting that these may be the Zn^2+^ sensing fingers of the protein. Even though most TFs that bind Zn^2+^ have K_D_s in the hundreds of picomolar range, they are generally not thought to be metal responsive. For example, it’s been shown that the nuclear hormone receptors glucocorticoid receptor (GR) and estrogen receptor (ER) contain two C4 Zn^2+^ fingers bind Zn^2+^ with K_D_s of 316 and 501 pM, respectively^21^. Given that it has been shown that cells experience changes in the labile Zn^2+^ pool^22–27^, it is important to establish whether other Zn^2+^ finger TFs are susceptible to changes in the Zn^2+^ pool.

In this work, we sought to determine whether perturbations in the labile Zn^2+^ pool result in changes to TF activities. We utilized fluorescently tagged versions of the GR, with two C4 Zn^2+^ finger domains, and CCCTC-binding factor (CTCF), with eleven C2H2 Zn^2+^ finger domains, and single molecule fluorescence microscopy to monitor their mobility within live cells. We found that CTCF, but not GR, shows increases in its displacement and apparent diffusion coefficient when cellular Zn^2+^ is chelated with Tris(2-pyridylmethyl)amine (TPA), suggesting that CTCF is more dynamic when cellular Zn^2+^ is low. However, when the dwell times for CTCF are calculated, we see that both chelation of Zn^2+^ and addition of Zn^2+^ results in a reduction in the dwell time of CTCF. This could point to larger changes in chromatin architecture that occur when Zn^2+^ is perturbed.

## Results

While it is well established that Zn^2+^ finger transcription factors require Zn^2+^ to bind DNA in vitro, an open question is whether the Zn^2+^ occupancy and hence DNA binding capacity in cells is dependent on levels of cellular Zn^2+^. We applied 3D single molecule microscopy to investigate whether Zn^2+^ finger transcription factors have altered mobility when Zn^2+^ is perturbed, where mobility is routinely used as a proxy for DNA binding^29,33–36^. Specifically, we used multifocus microscopy (MFM) that allows for simultaneous acquisition of particles in 9 Z-planes separated by approximately 430 nm (~ 3.9 μm total axial depth)^30^. This enabled us to track labeled transcription factors with high accuracy at relatively rapid acquisition rates (25 Hz) and did not result in truncated trajectories as they diffused along the Z-axis in the nucleus, a bottleneck that can be seen with 2D single molecule tracking.

For candidate transcription factors, we selected the glucocorticoid receptor (GR), a nuclear hormone receptor containing two C4 Zn^2+^ finger domains, and the CCTCC-binding factor (CTCF), a chromatin binding protein that contains eleven C2H2 Zn^2+^ finger domains. Both transcription factors have been previously characterized by 2D single molecule microscopy and have DNA dwell times that span an approximate order of magnitude (3-8 s for GR^34^, 60 s for CTCF^29^). U2OS cells expressing HaloTag-GR (stable overexpression) or HaloTag-CTCF (CRISPR-edited endogenous expression) were treated with either 50 μM of the Zn^2+^ chelator TPA to deplete free Zn^2+^, 30 μM ZnCl_2_ to increase free Zn^2+^, or a media-only control for 30 mins. Previously, we have shown that these perturbations decrease labile Zn^2+^ to <1 pM or increase labile Zn^2+^ to 30 nM, respectively^22,23,37^. Under our imaging conditions, we were able to acquire between 493-1080 total tracks for HaloTag-GR and between 5065-12770 tracks for HaloTag-CTCF (Table 1). Overlaying 100 random trajectories for the Zn^2+^ replete and Zn^2+^ deficient conditions compared to control revealed apparent mobility differences for both HaloTag-CTCF and Halo-Tag-GR (Figure 1). Both TFs appeared to be more diffusive in Zn^2+^ deficiency compared to control media, in which the protein appeared to be more immobilized. Surprisingly, both TFs also appeared to be slightly more diffusive in Zn^2+^ rich media compared to the control.

**Figure 1.**
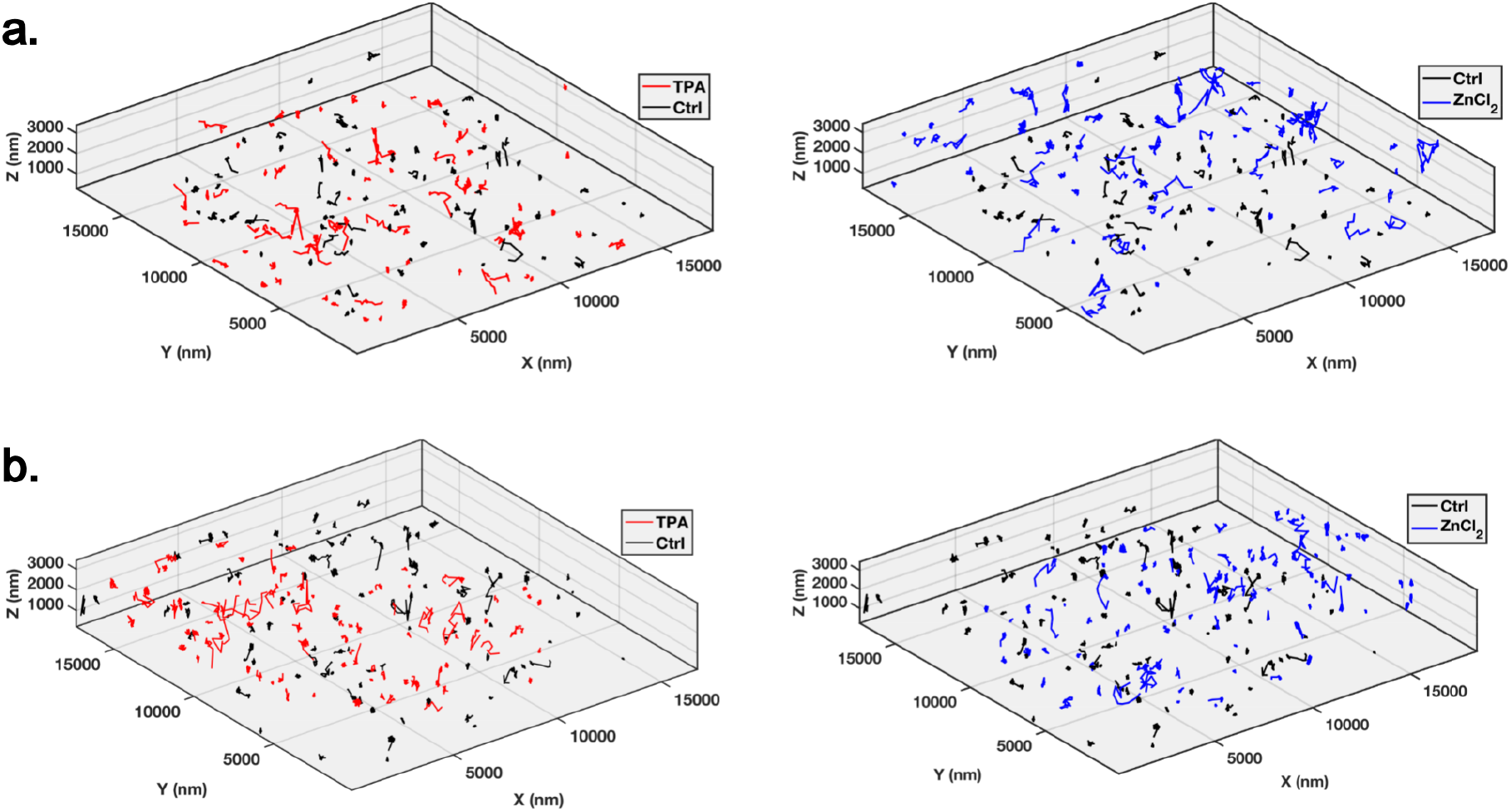
3D single molecule trajectories of HaloTag TFs. (A) HaloTag-GR cells were treated with either 50 μM TPA (left), 30 μM ZnCl_2_ (right), or a media-only control (left and right) for 30 mins. 100 tracks from one cell of each condition were overlaid on top of each other. (B) Same as in (A), but with HaloTag-CTCF cells. Total number of cells acquired and tracks detected per cell can be found in Table 1.

**Table 1.** Summary statistics for all cell lines and imaging conditions used in this study. Here, MFM denotes experiments conducted to generate 3D particle tracking data and N-STORM denotes experiments conducted to generate 2D partice tracking data.

| Cell Line | Treatment | Microscope | Exposure Time (ms) | Acquisition Interval (ms) | Number of Cells | Total Number of Tracks | Total Number of Tracks – Diffusion | Total Number of Dwell Times |
| --- | --- | --- | --- | --- | --- | --- | --- | --- |
| U2OS HaloTag-GR | 50 μM TPA | MFM | 40 | 40 | 5 | 493 | 262 | 385 |
| U2OS HaloTag-GR | Ctrl | MFM | 40 | 40 | 6 | 948 | 419 | 630 |
| U2OS HaloTag-GR | 30 μM ZnCl<sub>2</sub> | MFM | 40 | 40 | 5 | 1080 | 601 | 708 |
| U2OS HaloTag-CTCF | 50 μM TPA | MFM | 40 | 40 | 11 | 11704 | 5046 | 12771 |
| U2OS HaloTag-CTCF | Ctrl | MFM | 40 | 40 | 10 | 12770 | 4985 | 14295 |
| U2OS HaloTag-CTCF | 30 μM ZnCl<sub>2</sub> | MFM | 40 | 40 | 7 | 5065 | 1975 | 5958 |
| U2OS HaloTag-GR | 50 μM TPA | N-STORM | 100 | 100 | 8 | 17511 | N/A | 7702 |
| U2OS HaloTag-GR | Ctrl | N-STORM | 100 | 100 | 11 | 45748 | N/A | 10994 |
| U2OS HaloTag-GR | 30 μM ZnCl<sub>2</sub> | N-STORM | 100 | 100 | 9 | 24544 | N/A | 20121 |
| U2OS HaloTag-CTCF | 50 μM TPA | N-STORM | 100 | 500 | 5 | 12452 | N/A | 2721 |
| U2OS HaloTag-CTCF | Ctrl | N-STORM | 100 | 500 | 6 | 44628 | N/A | 12430 |
| U2OS HaloTag-CTCF | 30 μM ZnCl<sub>2</sub> | N-STORM | 100 | 500 | 7 | 21725 | N/A | 6179 |

While the single molecule trajectories suggested an apparent changes in mobility upon manipulation of Zn^2+^, we wanted to examine this in greater depth this via quantitative analysis. We first quantified this by plotting the cumulative distribution functions (CDF) and survival curves (1-CDF) of the displacements for the first 8 frames of each trajectory (Figure 2A). For each time delay (Δt_1_: 0-40 ms; Δt_2_: 0-80 ms; Δt_3_: 0-120 ms, etc.), we calculated how far each TF traveled and plotted the survival curves for each time delay. Overall, the displacement increased with increasing time delay (Δt_1_ to Δt_4_ to Δt_8_), as expected. At all time delays, HaloTag-GR appears to be more mobile with both TPA and Zn2Cl treatment relative to the control, although the increase in displacement with TPA or Zn^2+^ is small (Figures 2B and 2D). In contrast, HaloTag-CTCF appears to be more mobile only upon treatment with TPA relative to the control (Figures 2C and 2E). The increase in displacement upon treatment with TPA is largest for the highest time delay (Δt_8_). Furthermore, the increase in displacement with TPA is much higher in magnitude compared to GR.

**Figure 2.**
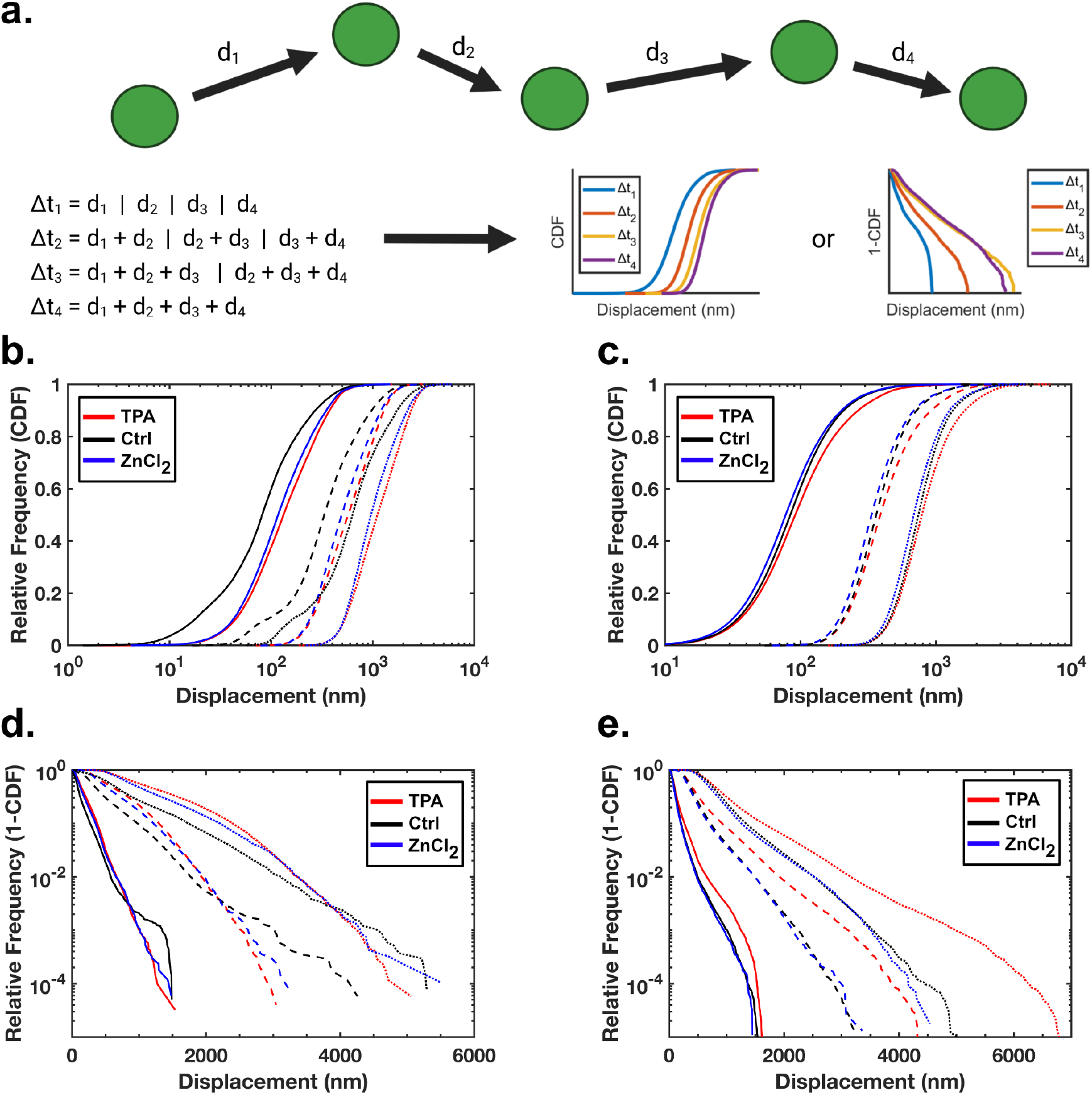
(A) Schematic showing how particle displacements are calculated. For each particle, the distances traveled during each time delay (Δt_1_ (40 ms), Δt_2_ (80 ms), etc.) were compiled and both the cumulative distribution functions (CDF) and survival curves (1-CDF) for each time delay were subsequently plotted. (B) Displacement CDFs for HaloTag-GR cells treated with either 50 μM TPA (red), a media control (black), or 30 μM ZnCl_2_ (blue) for 30 min. Data for each treatment are shown for Δt_1_ (40 ms; solid lines), Δt_4_ (160 ms; dashed lines), and Δt_8_ (320 ms; dashed lines). (C) Displacement CDFs for HaloTag-CTCF cells. Coloring and line styles are the same as in (B). (D) Displacement survival curves for HaloTag-GR. Coloring and line styles are the same as in (B). (E) Displacement survival curves for HaloTag-CTCF. Coloring and line styles are the same as in (D).

To further quantify the mobility of HaloTag-GR and HaloTag-CTCF upon manipulating cellular Zn^2+^, we determined apparent diffusion coefficients. To determine the diffusion coefficient, particle trajectories were fit to an anomalous diffusion model *MSD* = *γDΔt^α^*, where *MSD* is the mean squared displacement, *γ* is the number of dimensions (here, 3) multiplied by 2, *D* is the apparent diffusion coefficient, *Δt* is the time delay, and *α* defines whether the particle is superdiffusive (*α* > 1) or subdiffusive (*α* < 1). While HaloTag-GR showed larger displacements with both TPA and ZnCl_2_ treatment, the calculated apparent diffusion coefficients did not vary significantly across conditions (Figure 3A). With HaloTag-CTCF, the apparent diffusion coefficient for TPA treatment is indeed significantly higher than both the control and ZnCl_2_-treated conditions (Figure 3B). This suggests that HaloTag-CTCF is indeed more mobile in Zn^2+^ deficient conditions, and that there is no difference between control and Zn^2+^ rich conditions.

**Figure 3.**
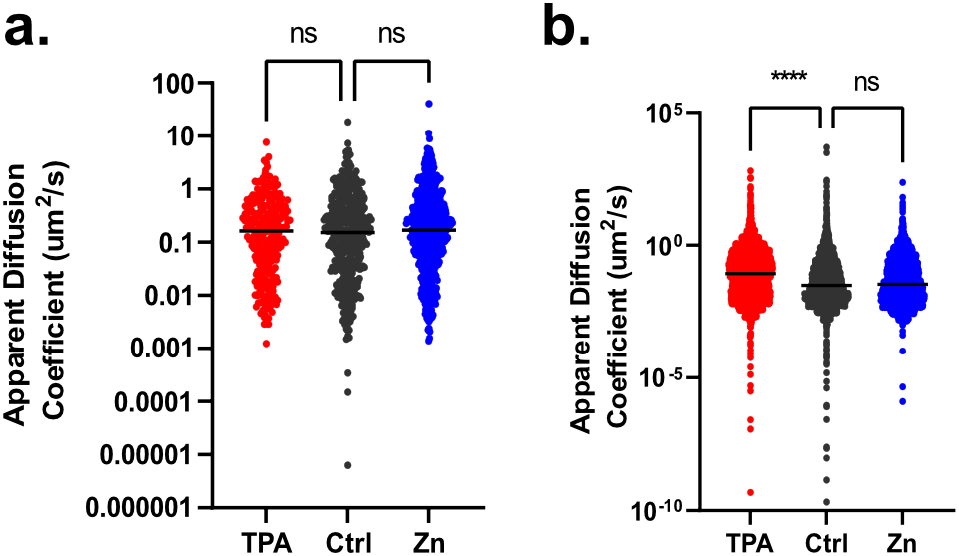
Apparent diffusion coefficients for (A) HaloTag-GR or (B) HaloTag-CTCF upon treatment with 50 μM TPA, 30 μM Zn, or a media-only Control. Each dot represents the apparent diffusion coefficient for a single TF trajectory. Plots represent > 260 TFs (for HaloTag-GR) and > 1975 TFs (for HaloTag-CTCF), from at least 5 cells for each measurement condition. For HaloTag-CTCF treatments, **** = p < 0.0001.

The above analysis suggests that CTCF shows increased displacement and greater diffusion upon Zn^2+^ depletion, while GR is minimally/not affected by Zn^2+^ depletion. To determine how long a particle (i.e. a TF) remains in one spot for a given amount of time, we computed the aggregate dwell times. Our rationale was that particles with longer dwell times may correspond to those that are bound to DNA rather than freely diffusing or experiencing short, transient interactions^29,33,34,36,38^. The displacements between each point of trajectories were calculated and consecutive displacements of less than 200 nm resulted in a particle being classified as bound. This 200 nm threshold has previously been used as a conservative estimate of general chromatin movement, as measured using fluorescently-labeled histone H2B^33,34^. Additionally, binding events had to last at least 8 frames (320 ms) to be considered bound; this eliminated transient, non-specific interactions that were sometimes observed.

To validate this method for calculating dwell times, we treated HaloTag-GR cells with the known activators dexamethasone or hydrocortisone. Previously it has been shown that dexamethasone is a more potent activator of GR than hydrocortisone and this leads to longer dwell times^34^, presumably because GR is more strongly associated with target sites on DNA. Consistent with previous reports, we found that dexamethasone shows a decreased apparent diffusion coefficient and an increased dwell time compared to hydrocortisone (Figure 4A and B). Having established that we could detect changes in dwell time from MFM datasets, we examined whether manipulation of Zn^2+^ altered the dwell time of GR and CTCF. Based on our other single molecule mobility measurements, we predicted that HaloTag-CTCF would show reduced dwell times upon treatment with TPA. However, neither HaloTag-GR nor HaloTag-CTCF show significantly different dwell times across treatment conditions. Surprisingly, the observed dwell times were substantially shorter than previous reports in the literature (less than 400 ms for both GR and CTCF vs ~8 sec for GR^34^ and ~ 60 sec for CTCF^29^) (Table 2).

**Figure 4.**
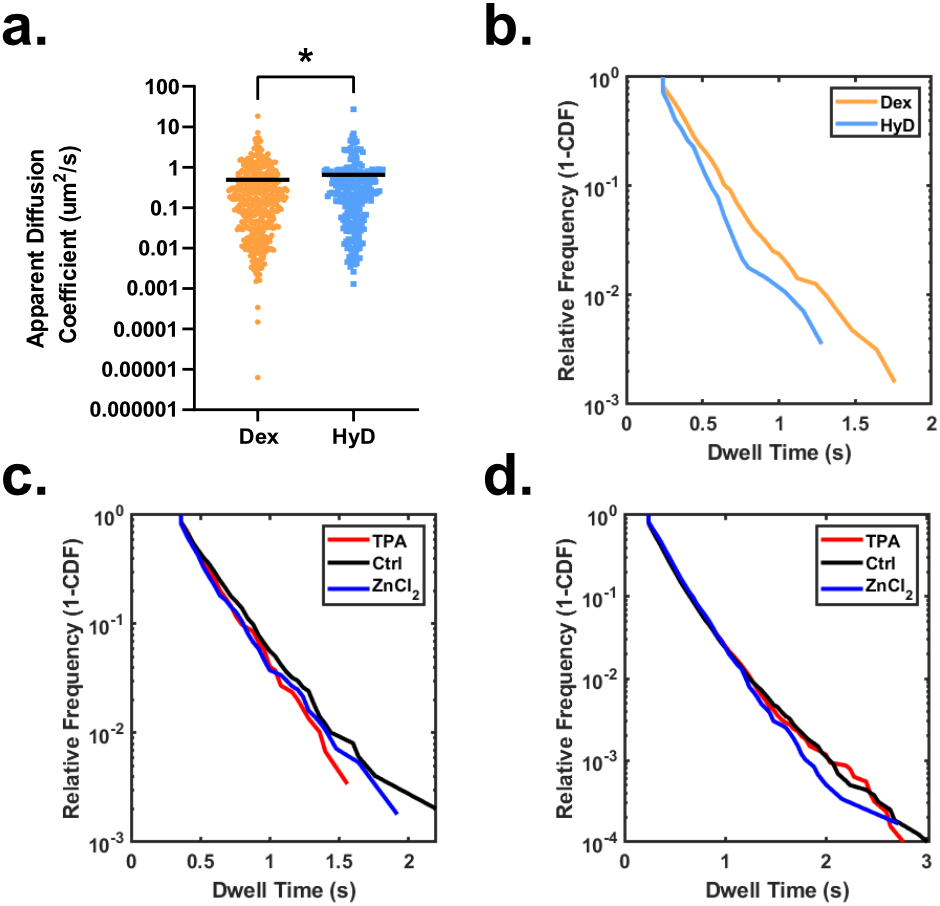
(A) Apparent diffusion coefficients for HaloTag-GR treated with either 100 nM dexamethasone (Dex) or 100 nM hydrocortisone (HyD). * denotes p = 0.01 (Mann-Whitney t-test). (B) Dwell time analysis for HaloTag-GR treated with either 100 nM dexamethasone or 100 nM hydrocortisone. (C) Dwell time analysis for HaloTag-GR treated with 50 μM TPA (red), 30 μM ZnCl_2_ (blue), or a media-only control (black). (D) Same as in (C) but with HaloTag-CTCF.

**Table 2.** 3D dwell time fits for HaloTag-GR and HaloTag-CTCF. For the two-component fit, *k1* and *t1* are the rate constant and dwell times for the fast component, and *k2* and *t2* are the rate constant and dwell times for the slow component.

|  |  | Two-Component Fit |  |  |  |
| --- | --- | --- | --- | --- | --- |
| Factor | Treatment | $k1$ | $t1$ | $k2$ | $t2$ |
| HaloTag-GR | 50 μM TPA | -5.98 | 0.167 | -134.50 | 0.01 |
| HaloTag-GR | Ctrl | -5.13 | 0.195 | -136.80 | 0.01 |
| HaloTag-GR | 30 μM ZnCl<sub>2</sub> | -106.90 | 0.009 | -6.27 | 0.16 |
| HaloTag-CTCF | 50 μM TPA | -7.56 | 0.132 | -3.39 | 0.29 |
| HaloTag-CTCF | Ctrl | -5.99 | 0.167 | -2.74 | 0.37 |
| HaloTag-CTCF | 30 μM ZnCl<sub>2</sub> | -135.30 | 0.007 | -4.91 | 0.20 |

We hypothesized that perhaps our measurement window on the MFM was too short to accurately determine the dwell time for CTCF. This measurement window was limited to 2 min of tracking data due to photobleaching of the JF dye under the illumination conditions of the MFM. Indeed, the literature measurement of CTCF dwell time was approximately 1 min^29^, suggesting a 2 min measurement period would be insufficient for accurately determining the dwell time. Therefore, we performed a similar experiment but instead used a Nikon N-STORM microscope in highly-inclined laminated optical sheet (HILO)^39^ mode with longer acquisition times and substantially less laser intensity. Imaging using HILO allowed for the laser to illuminate particles within the nucleus and by greatly reducing the intensity of the illumination laser, we were able to image for 5 mins at 10 Hz (for HaloTag-GR) and 20-30 mins at 2 Hz (for HaloTag-CTCF) rather than the 1-2 min acquisition periods on the MFM. Additionally, these conditions allowed us to bias all detected particles towards bound particles, rather than freely diffusing particles. As such, we calculated the particle dwell time to be equal to *track length × frame rate* (*s*), with a minimum track length of 5 to be considered in the analysis.

For HaloTag-GR, treatment with TPA or ZnCl_2_ did not significantly affect particle dwell times, consistent with other measurements showing that perturbation of Zn^2+^ did not significantly alter mobility of HaloTag-GR. For HaloTag-CTCF, TPA caused a reduction in particle dwell times compared to the control untreated cells. This correlates with the 3D particle tracking and diffusion data we measured on the MFM and suggests that Zn^2+^ depletion indeed causes greater mobility for CTCF. Treatment with ZnCl_2_ also led to a slight reduction in observed dwell times compared to the untreated control, although the magnitude of the shift was not as much as TPA treatment. The measured dwell times were consistent with previously measured dwell times^29,34^ (Table 3).

**Table 3.** Photobleaching-corrected 2D dwell time fits for HaloTag-GR and HaloTag-CTCF. *k1* and *t1* are the rate constant and dwell times for the fast component, and *k2* and *t2* are the rate constant and dwell times for the slow component.

|  |  | Two-Component Exponential Fit |  |  |  |
| --- | --- | --- | --- | --- | --- |
| Factor | Treatment | $k1$ | $t1$ | $k2$ | $t2$ |
| HaloTag-GR | 50 μM TPA | -0.89 | 1.13 | -0.13 | 7.48 |
| HaloTag-GR | Ctrl | -0.10 | 10.03 | -0.58 | 1.73 |
| HaloTag-GR | 30 μM ZnCl<sub>2</sub> | -0.91 | 1.10 | -0.15 | 6.73 |
| HaloTag-CTCF | 50 μM TPA | -0.03 | 29.35 | -0.24 | 4.12 |
| HaloTag-CTCF | Ctrl | -0.13 | 7.91 | -0.03 | 35.48 |
| HaloTag-CTCF | 30 μM ZnCl<sub>2</sub> | -0.22 | 4.56 | -0.04 | 22.46 |

Previous studies showed that deletion of the 11 Zn^2+^ fingers in CTCF resulted in long displacements consistent with free diffusion of CTCF^29^. This suggests that without Zn^2+^ fingers, CTCF is unable to effectively bind DNA. We set out to determine whether depletion of Zn^2+^ can alter the bound versus unbound trajectories for both HaloTag-GR and HaloTag-CTCF. The fraction bound was determined by dividing the number of trajectories longer than 5 frames by the total number of trajectories. While our imaging conditions did bias detections towards bound tracks, we only computed dwell times for trajectories lasting longer than 5 frames. For HaloTag-GR, we found that the fraction bound did not vary significantly across treatments (Figure 5C). For HaloTag-CTCF, we found that the fraction bound for TPA treated cells was less than the control and ZnCl_2_ treated cells, with a p value = 0.0572 (Figure 5D). We didn’t observe any difference between the cells treated with ZnCl_2_ and the untreated control.

**Figure 5.**
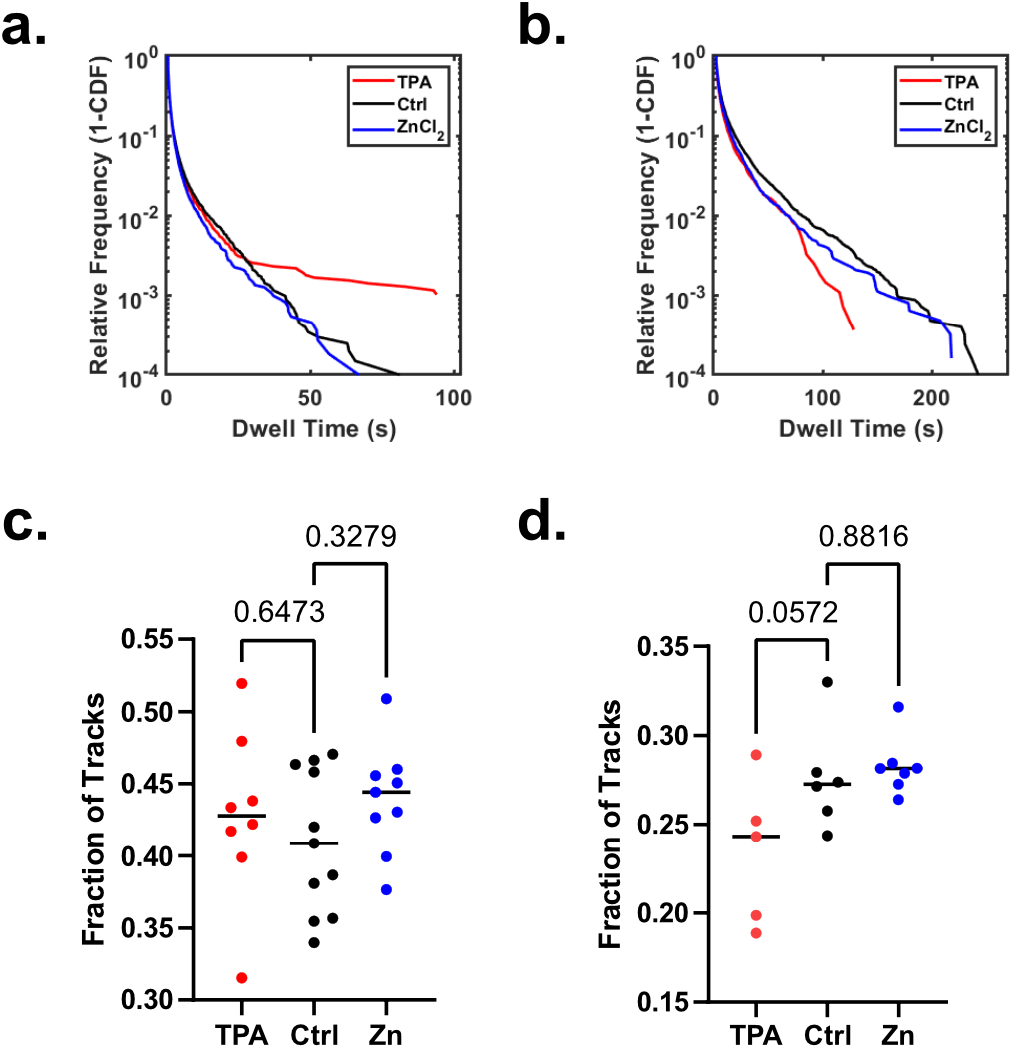
(A) 2D dwell time survival curves (1-CDF) for HaloTag-GR cells treated with 50 μM TPA (red), 30 μM ZnCl_2_ (blue), or a media-only control (black). (B) same as in (A), but with HaloTag-CTCF cells. (C) Fraction bound analysis for HaloTag-GR cells. Fraction bound was calculated as the number of tracks that were present for at least 5 frames (500 ms) divided by the total number of detected tracks. (D) Same as in (C), except with HaloTag-CTCF cells and with tracks that were present for at least 5 frames (2.5 s). For (C) and (D), numbers indicate the p-value calculated using a one-way ANOVA.

## Discussion

Proper transcriptional regulation is essential for the cell’s ability to respond to the demands of its environment. Over the past several decades, various genomic techniques have been developed to monitor TF activity and the downstream consequences on gene expression, but most of these techniques monitor a heterogenous population of cells at fixed time points. SM microscopy allows for assessment of individual TFs within single cells and can easily be scaled to accommodate as many treatment time points as needed. However, it is important to keep in mind precisely which measurements are desired at the end of the SM experiment, as this will dictate the imaging setup and acquisition parameters needed. For example, the MFM used in this study was designed with the goal of enabling simultaneous rapid acquisition of multiple Z-planes. Additionally, many standard microscopes that are capable of total internal reflection fluorescent (TIRF) microscopy can be modulated to perform HILO imaging. When acquiring at fast (>25 Hz) frame rates, both systems are capable of measuring “fast” TF characteristics such as diffusion coefficients. However, under these illumination conditions even the most robust fluorophores will rapidly bleach, resulting in relatively short trajectories. Indeed, this was seen with both HaloTag-GR and HaloTag-CTCF within our MFM data: almost no trajectories existed for more than 3 seconds (Figure 4). By implementing HILO imaging with less intense laser power and a longer camera acquisition time, we were able to measure much longer trajectories (in rare cases, up to 4 mins for HaloTag-CTCF) and calculate dwell times that were on the time scales previously reported in the literature.^29,34^

It is hypothesized that up to 10% of the human proteome encodes for Zn^2+^ binding proteins, and nearly half (44%) are involved in transcriptional regulation^40,41^. However, very little is known about the metal status of these proteins *in vivo*. A small subset of Zn^2+^ binding TFs has K_D_s reported in the literature, but these tend to be measured in the context of a single Zn^2+^ finger as opposed to the entire protein^17^. It is unknown how they are metalated and how the proteins respond to changes in the labile Zn^2+^ pool. Mass spectrometry techniques have been developed that seek to label Zn^2+^-binding cysteine residues^42^ but so far this has only been done in the context of addition of Zn^2+^ or EDTA to cell lysates^42^. Additionally, TFs tend to exist in relatively low abundance compared to cytosolic proteins, so special experimental considerations must be taken to enrich for potential targets.

In this work, we found that the dynamics of HaloTag-GR and HaloTag-CTCF are both susceptible to changes in cellular Zn^2+^ status but in subtly different ways. When examining the overall displacements of HaloTag-GR, it appears that treatment with either TPA or ZnCl_2_ results in more mobile proteins. However, this does not translate to any measurable change in diffusion coefficient. It could be that we employed an overly simplistic model of diffusion and that newer models^43^ may be more robust but these are mostly designed for 2D datasets. In contrast, HaloTag-CTCF both appeared to be more dynamic upon TPA treatment and had an increased diffusion coefficient, indicating that in Zn^2+^ deficiency it may be more mobile. When characterizing HaloTag-CTCF’s dwell times via 2D tracking, we found that both TPA and ZnCl_2_ caused a reduction in dwell times.

The differences in mobility, diffusion, and dwell time between GR and CTCF that we present here can be rationalized in a few different ways. First, the number and type of Zn^2+^ fingers used by these two proteins differ, with CTCF having 11 C2H2 Zn^2+^ fingers and GR having 2 C4 Zn^2+^ fingers. While the literature has shown that the mode of Zn^2+^ binding does not necessarily dictate the relative affinity of the Zn^2+^ ion for isolated Zn^2+^ fingers^17^, the binding mode may come more into play when looking at full length proteins versus isolated DNA binding domains due to the larger second coordination sphere. Additionally, CTCF has been shown to utilize different combinations of its Zn^2+^ fingers to bind distinct DNA sequences, allowing it to bind 80,000+ sites in the human genome^44^. While there is likely some redundancy in which of CTCF’s Zn^2+^ fingers are binding to DNA, it is plausible that perturbing the Zn^2+^ status of even one of these domains may affect the ability of CTCF to adequately bind its genomic targets. Further, it is important to note that the dwell times we observe do not necessarily indicate functional binding events. As noted, CTCF can potentially bind 80,000+ sites in the genome, and GR ChIP-seq experiments have shown 10,000+ potential binding sites across multiple cell types^45^. However, of the ^~^7000 sites found in A549 (lung adenocarcinoma) cells, only 928 (13.5%) of these sites were shown to truly be glucocorticoid responsive. It is therefore plausible that some of the binding events we observe via single molecule microscopy may be dominated by transient, non-specific interactions rather than productive binding. This is where newer quantitative methods^46^ for assessing particle dwell time will excel and help the field deconvolve the complexity of transcriptional regulation.

Our work suggests that a subset of TFs may be able to sense changes in cellular Zn^2+^. What this means from a functional standpoint remains to be seen. Given the noted advantages of SM microscopy, it would be valuable to pair these techniques with other fluorescent microscopy techniques to assess the downstream consequences of this sensing. GR, as a canonical TF, could be paired with the incorporation of promoter arrays to more readily assess specific binding^34^, or it could be coupled with nascent RNA imaging^14,15^ to examine the consequences on its target gene products. CTCF, as a regulator of chromatin architecture, could be paired with microscopy techniques that measure chromatin compaction^47–49^ to see if this is perturbed with changes in Zn^2+^. These tools will better allow us to understand the critical role that Zn^2+^ homeostasis fulfills in the cell.

## Materials & Methods

### Plasmid Generation

To generate PiggyBac-GR-HaloTag, PB-CMV-MCS-EF1α-Puro (System Bioscience #PB510B-1) was linearized using EcoRI and BamHI. The GR insert was amplified from pk7-GR-GFP (Addgene #15534) to generate overhangs with both PB-CMV-MCS-EF1α-Puro and the HaloTag. The HaloTag insert was amplified from pcDNA3.1-3xFLAG-HaloTag-2xNLS (Daniel Youmans, Cech lab, CU Boulder) using the primers listed in the Key Resources table to generate overhangs with GR and PB-CMV-MCS-EF1α-Puro. Linear fragments were then assembled into the final plasmid using a homemade Gibson assembly master mix.^28^

### Cell Culture Conditions

All cells were cultured in Dulbecco’s Modified Eagle Medium (DMEM) (ThermoFisher #12800082) containing 10% FBS (SigmaAldrich #F0926)) and 1% penicillin/streptomycin (ThermoFisher #15140-200). All live cell single molecule tracking experiments were carried out in FluoroBrite DMEM (ThermoFisher #A1896701). To generate stable U-2 OS HaloTag-GR cells, wildtype U-2 OS cells (ATCC #HTB-96) were transfected with 1000 μg of PiggyBac-HaloTag GR and 250 μg of Super PiggyBac Transposase (System BioSciences #PB200A-1) and 3 μL of TransIt LT1 (Mirus #MIR2305). Stable clones were selected by growing in DMEM containing 0.5 μg/μL puromycin (Sigma-Aldrich #P8833-25MG) for 7 days, after which they were transferred to normal DMEM. Cells expressing endogenous HaloTag-CTCF were a gift from Anders Serj Hansen (MIT)^29^.

### Zinc Perturbations

To manipulate labile Zn^2+^, cells were treated with either 30 μM ZnCl_2_ (Sigma-Aldrich # 39059-100ML-F) or 50 μM Tris(2-pyridylmethyl)amine (TPA, Sigma-Aldrich # 723134-250MG) for 30 mins prior to imaging. For experiments involving HaloTag-GR, Zn^2+^ perturbations were followed by hormone activation with either 100 nM dexamethasone (Sigma-Aldrich #D4902-100MG) or 100 nM hydrocortisone (Sigma-Aldrich #H4001-1G)

### 3D Single Particle Tracking

Three dimensional single particle tracking was performed on the Multifocus Microscope (MFM) at the Janelia Advanced Imaging Center (HHMI)^30^. Briefly, an epi-fluorescent microscope equipped with a multi-focal diffraction grating (MFG) allows for the collection of 9 aberration-corrected focal planes. Chromatic aberrations are corrected using a separate chromatic correction grating and prism. The MFG allows for a total axial detection depth of approximately 4 μm. For each experimental day, a calibration with TetraSpeck 0.2 um fluorescent beads (ThermoFisher #T7279) was used to determine the precise Z-spacing between each focal plane, with an average ΔZ of 430 nm. Additionally, the calibration allowed for measurement of the point spread function of the microscope, which allowed for image deconvolution (see below). All images were acquired using a 100x 1.45 NA TIRF objective (Nikon), a 561 nm laser (Cobolt Jive 300,), a Di01-R405/488/561/635 dichroic (Semrock), a FF01-593/40 (Semrock) emission filter, and an iXon3-DU897E EMCCD (Andor Technologies).

#### Image Acquisition

Cells were labeled with 1 μM of JaneliaFluor (JF) 549 HaloTag ligand (Janelia Research Campus) for 5 min at 37°C, rinsed three times with Dulbecco’s phosphate-buffered saline (D-PBS), and then incubated for 30 mins in FluoroBrite DMEM. Image acquisition was performed using a 561 nm laser at typical irradiance of 3-4kW/cm^2^, with 40 ms exposure times for an effective frame rate of 25 Hz. Movies, on average, were acquired for 2 mins (3000 frames).

#### Post Processing

Following acquisition, movies were cropped and laterally registered using a pre-determined affine transformation determined via TetraSpeck bead data (described above) to convert the 3×3 image into a 9 Z-plane stack, spanning approximately 3.8 μm in the Z-dimension. To improve signal-to-noise ratio and particle localization, local background was subtracted using a rolling ball correction (radius = 7 px), followed by 5 iterations of the Richardson-Lucy deconvolution algorithm within MATLAB R2019a (Mathworks). Following deconvolution, images were smoothed using a Gaussian filter (radius = 0.7 px) to improve particle detection. Additionally, the first 500 frames of each movie were removed, as these tended to have dense labeling that did not allow for robust tracking.

#### 3D Particle Tracking

Particle trajectories were generated using the MosaicSuite ImageJ plugin^31^ with the following parameters: radius = 3 px; cutoff = 0.001; threshold = 750; max link range = 1 frame; max displacement = 500 nm; dynamics = Brownian. Additionally, any tracks that did not exist for at least 10 frames (400 ms) were discarded.

#### Diffusion analysis

To determine diffusion coefficients for HaloTag-GR and HaloTag-CTCF, the mean squared displacement (MSD) for each track was produced and this was fit to the equation *MSD* = *γDΔt^α^*, where *MSD* is the mean squared displacement, *γ* is the number of dimensions (3) multiplied by 2, *D* is the apparent diffusion coefficient, *Δt* is the time delay between frames (here, 40 ms), and *α* defines whether the particle is superdiffusive (*α* > 1) or subdiffusive (*α* < 1). Because transcription factors are confined within the nucleus, they tend to exhibit anomalous diffusion (alpha < 1), rather than Brownian motion (alpha = 1) or superdiffusion (alpha > 1)^32^. For this analysis, tracks which did not fit an anomalous model were discarded, as were tracks that did not yield a good fit (R^2^ < 0.6).

#### Particle dwell time analysis

To compute 3D particle dwell times, cumulative displacements from one frame to each subsequent frame were measured. If the displacement from one frame through the next 5 frames was less than a pre-determined threshold distance (300 nm, or the maximum distance a bound molecule of H2B is known to move at these frame rates^33^), then the particle would be considered bound. This calculation was performed for each portion of the trajectory, so if a binding event occurred later in the track rather than the beginning, it would still be captured.

### 2D Single Particle Tracking

#### Equipment

2D single particle images were acquired on a Nikon N-STORM imaging system equipped with a Nikon TI-E microscope, a Nikon CFI Apo TIRF 100X oil immersion objective (1.49 NA), a N-STORM 647 nm laser (Agilent), an iXon 897 Ultra EMCCD (Andor Technologies), and a cage incubator (Okolab).

#### Image Acquisition

Cells were stained with JaneliaFluor (JF) 646 HaloTag ligand at either 100 pM (U-2 OS HaloTag-CTCF) for 1 min or 1 nM (U-2 OS HaloTag-GR) for 5 min, rinsed three times with D-PBS, and then incubated for 30 mins in FluoroBrite DMEM prior to Zn^2+^ perturbations. Image acquisition occurred with low laser intensities (5-10%) and 100 ms exposure times. Frame rates were chosen to bias towards only detecting bound particles: for HaloTag-CTCF, effective frame rate = 2 Hz; for HaloTag-GR, effective frame rate = 10 Hz. Because CTCF is known to have long residence times on DNA, these movies were collected on average for 20 mins (2400 frames), while movies for GR were typically acquired for 2-5 mins.

#### Post processing

Movies were post-processed within Nikon Elements to subtract background using the rolling ball method (radius = 50), and then subjected to 5 iterations of the Richardson-Lucy deconvolution algorithm within Nikon Elements. Following deconvolution, images were smoothed using a Gaussian filter (radius = 0.7 px) to improve particle detection. Additionally, the first 500 frames of each movie were excluded, as these tended to have dense labeling that did not allow for robust tracking.

#### 2D Particle Tracking

Particle trajectories were generated using the MosaicSuite ImageJ plugin with the following parameters: radius = 3 px; cutoff = 0; absolute threshold = 500-1000, depending on the experiment; max link range = 1 frame; max displacement = 300 nm; dynamics = Brownian. Additionally, any tracks that did not exist for at least 5 frames (2.5 sec for CTCF and H2B; 0.5 sec for HaloTag-GR) were excluded from further analysis.

#### Dwell time analysis

Because the effective frame rate of each movie and tracking parameters were biased towards only detecting bound molecules, we inferred that the only particles detected were bound. Therefore, we calculated dwell time as the length of the track divided by the frame rate, and the aggregate dwell times were used to generate dwell time survival curves. These were subsequently corrected for photobleaching as previously described^33,34^. Briefly, the rate of photobleaching was estimated by fitting the detected number of particles per frame and fitting this to the two-component exponential decay, *B*(*t*) = *ae*^(*k*_*b*1_*t*)^ + *be*^(*k*_*b*2_*t*)^, where *a* and *b* are the fraction sizes of the two components and *k*_*b*1_ and *k*_*b*2_ are the photobleaching rates. The dwell time survival curves were then divided by *B*(*t*) and subsequently fit using the equation *f*(*t*) = *ae*^(*k*_1_*t*)^ + *be*^(*k*_2_*t*)^, where *a* and *b* are the fraction sizes of the two components and *k*_1_ and *k*_2_ are the rate constants. Dwell times (*τ*) for each component were subsequently computed as 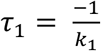 and 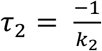.

## Acknowledgements

We would like the thank the following sources for financial support: NIH Director’s Pioneer Award DP1-GM114863 (AEP), NIGMS MIRA R35 GM139644 (AEP), Molecular Biophysics Training Grant T32 GM-065103 (LJD). Aberration corrected multifocus microscopy was performed in collaboration with the Advanced Imaging Center at Janelia Research Campus, a facility jointly supported by the Gordon and Betty Moore Foundation and the Howard Hughes Medical Institute. We would like to acknowledge the BioFrontiers Institute Advanced Light Microscopy Core and Dr. Joseph Dragavon, supported by the BioFrontiers Institute and the Howard Hughes Medical Institute. We would also like to acknowledge the University of Colorado Biochemistry Cell Culture Core Facility, especially Theresa Nahreini, for providing resources and support for all our cell work. We would like to thank Nick Lammer for helpful discussion and insight on glucocorticoid receptor. We thank Anders Serj Hansen (MIT) for generous gift of HaloTag-CTCF U-2 OS cells.

## Author contributions

LJD and AEP conceived of the study. LJD carried out experiments. JA assisted with multifocus microscopy data collection, processing and analysis. LJD and AEP analyzed data and wrote the paper. JA edited the paper.

## Data Availability statement

Time-lapse imaging data available on request.

